# Estimating descending activation patterns from EMG in fast and slow movements using a model of the stretch reflex

**DOI:** 10.1101/2024.10.01.615748

**Authors:** Lei Zhang, Gregor Schöner

## Abstract

Due to spinal reflex loops, descending activation from the brain is not the only source of muscle activation that ultimately generates movement. This study directly estimates descending activation patterns from measured patterns of muscle activation (EMG) during human arm movements. A simple model of the spinal stretch reflex is calibrated in a postural unloading task and then used to estimate descending activation patterns from muscle EMG patterns and kinematics during voluntary arm motion performed at different speeds. We observed three key features of the estimated descending activation patterns: (1) Within about the first 15% of movement duration, descending and muscle activations are temporally aligned. Thereafter, they diverge and develop qualitatively different temporal profiles. (2) The time course of descending activation is monotonic for slow movements, non-monotonic for fast movements. (3) Varying model parameters like the spinal reflex gain or the level of co-contraction does not qualitatively change the temporal pattern of estimated descending activation. Our findings highlight the substantial contribution of spinal reflex loops to movement generation, while at the same time providing evidence that the brain must generate qualitatively different descending activation patterns for movements that vary in their mechanical dynamics.

**New & Noteworthy:** We propose a new method that directly estimates descending activation from measured EMG signals and arm kinematics by inverting a model of the spinal stretch reflex, without the need for muscle models or for an arm dynamics model. This approach identifies key features of the time structure of descending activation as movement speed is varied, while also revealing the significant contribution of the spinal stretch reflex to movement generation.

## Introduction

The spinal stretch reflex contributes to the production of muscle forces and plays a significant role in movement generation (Nielsen 2016). Unlike the long-latency transcortical reflex, which operates with a relatively long delay and provides flexible modulation (Lee and Tatton 1982, Pruszynski and Scott 2012), the spinal stretch reflex is characterized by its short latency and is often perceived as a more stereotyped response (Pierrot-Deseilligny and Burke 2005). There is ample evidence supporting the important role of the spinal stretch reflex in the generation of voluntary movements. Mechanical perturbations can elicit this reflex (Bennett 1993, Smeets et al. 1995, Adamovich et al. 1997, Shapiro et al. 2004), and spinal afferent neurons maintain activity in force-producing tasks, as shown by direct recordings (Edin and Vallbo 1990, Dimitriou 2022, Umeda et al. 2022) or peripheral nerve stimulations (Janswoka 1992, Hultborn 2006, Nielsen 2016). Despite this role, spinal reflex loops are not taken into account in many contemporary theories of voluntary movement (Todorov and Jordan 2002, Churchland and Shenoy 2024). This suggests a need to characterize more precisely how the spinal stretch reflex contributes to muscle activation and movement generation.

Model simulations suggest that spinal reflexes may allow for relatively simple patterns of descending activation to bring about movements (Gribble et al. 1998, Pilon and Feldman 2006, Raphael et al. 2010, Buhrmann and Paolo 2014, Zhang et al. 2016, Niyo et al. 2024). These model simulations made use of temporal patterns of descending activation that were assumed ad-hoc, typically in the form of activation ramps or impulses. Estimating the descending activation pattern that is required to bring about a given movement requires some form of model inversion in which the movement trajectory is provided as input and the descending activation pattern is obtained as output of the inverted model. One attempt at such inversion combined a model of the spinal stretch reflex with a muscle model and a biomechanical model of the arm’s dynamics. Descending activation patterns that take the arm from an initial to a target position within a given movement time were estimated by minimizing the change of descending activation needed to move from the initial to the final posture (Ramadan et al. 2022). The minimally changing descending activation patterns found in that study had qualitatively different time structure when movement speeds were varied: slow movements were explained by monotonic, ramp-like descending activation patterns, fast movements by non-monotonic “N-shaped” descending activation patterns. Similar patterns of estimated descending activation were observed in a second study in which the same model was inverted analytically making a number of approximations. This made it possible to take experimentally measured joint kinematics to predict the time courses of descending activation (Hummert et al. 2024). In both studies, models of muscle dynamics and of the arm’s dynamics with considerable numbers of model parameters were required to estimate the descending activation patterns.

Here we propose a new method that estimates descending activation directly from measured EMG signals and arm kinematics and does not require muscle models or a model of the arm’s dynamics. The core idea is to invert a model of the spinal stretch reflex (Fig. 1) in which muscle (or motoneuron) activation *A* is composed of descending activation *u*_*d*_ and reflex activation *u*_*a*_.

**Figure 1.**
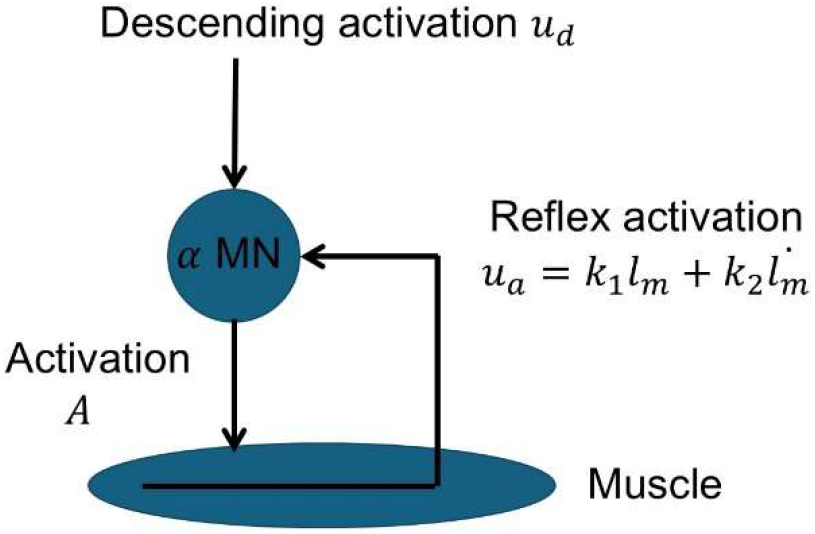
A reflex model in which muscle or motoneuron activation *A* depends on descending activation *u*_*d*_ and length and velocity-dependent reflex activation *ua*.

Muscle activation is the output of a semi-linear threshold function, with the intrinsic motoneuron threshold *u*_*Th*_. The threshold function [ ]^+^is equal to its input when the input value is not less than 0, otherwise it is 0:

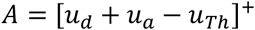

We further assume that the reflex activation depends linearly on muscle length *l*_*m*_ and velocity 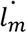:

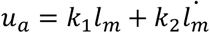

The reflex model is used to link muscle activation, *A*, to measured *EMG* in an arm postural task in which descending activation, *u*_*d*_, is assumed to remain unchanged when posture is shifted by removing an external load. In a voluntary movement task, muscle activation *A* is obtained from measured *EMG*, and muscle length, *l*_*m*_, and velocity, 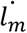, is obtained from measured joint angle *θ* and joint velocity 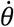, taking tendon properties into account. Descending activation is then estimated by analytically inverting the model, 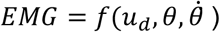, that results from these substitutions:

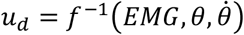

We estimate the temporal profile of *u*_*d*_ for voluntary arm movements at different speeds.

This estimation procedure enables us to address three important questions on the nature of descending movement control. First, how does the spinal stretch reflex contribute to movement generation? This will be revealed by comparing descending activation to muscle and reflex activation patterns. Second, how does descending control guide fast vs. slow movements? The key question here is whether the temporal forms of descending activation patterns driving fast movements are simply rescaled from those driving slow movements or whether they reflect the complex time structure of movement dynamics. Third, how do the temporal profiles of descending activation depend on certain independent control parameters, including the spinal feedback gain and co-contraction? This will be addressed by sensitivity analysis of these parameters in modulating the estimates of descending activation. We will discuss how our results relate to neurophysiological findings.

## Methods

### Participants

10 healthy participants (three males, age 31.0 ± 5.6 yrs, right-handed by self-reporting) participated in this study. All participants had no neurological disease that affects the upper extremity. They all signed an informed consent form approved by the institutional Ethics Committee in accordance with the 1964 Declaration of Helsinki.

### Experimental setup and data recording

Fig.2A and D shows the experimental setup for unloading and voluntary movements of the wrist. Participants sat comfortably in a dental chair that supported the head, neck and torso. The semi-supinated right forearm was fixed in a padded brace on a table (elbow angle ∼110 deg, shoulder horizontal abduction ∼45 deg). The hand with extended fingers was positioned in a plastic splint attached onto a light horizontal manipulandum. The hand was stabilized inside the splint filled with foam pads. The manipulandum could be rotated freely about a vertical axis aligned with the flexion–extension axis of the wrist joint, which the movement was restricted to. A torque motor (Parker Hannifin, Cleveland, OH, US) with maximal torque of 5 Nm was connected to the axis of the manipulandum and applied different levels of wrist flexion or extension torques (∼30% maximum voluntary contraction (MVC), range 0.4–1.2 N m) to the wrist. Unloading was produced by turning off the motor after preloading. During voluntary movement, the motor was off all the time.

The position of the manipulandum was recorded with an optical encoder coupled to the shaft of the manipulandum. Surface EMG activity was recorded from the flexor carpi radialis (FCR) and extensor carpi radialis (ECR) by a wireless EMG recording system (Delsys Trigno). Prior to electrode application, the skin was cleaned with alcohol. Electrodes were placed above the muscle bellies. A customized program (LabView, National Instruments, Austin, TX, USA) recorded data from the optical encoder of the torque motor and EMG signals (common sampling rate 2 kHz). The program also controlled the torque onset, duration, and magnitude of the torque motor. The current wrist position was displayed as a bar on a computer display in front of participants such that they could establish the initial wrist position in each trial. Neutral wrist position (when wrist muscles are fully relaxed) was defined as 0 deg. When the wrist flexes, the wrist angular position decreases. Positional feedback was unavailable after unloading. The initial position during unloading and the initial and end positions during voluntary movements were marked on the computer display.

### Experimental procedure

The main experiment consisted of two sessions. Session 1: unloading. Data of this postural task is used to estimate essential parameters in calibrating the neuromuscular model (which is explained in the last section of Methods). Session 2: voluntary movements. Data of this task will be used to reconstruct descending activations through the inversion of the calibrated neuromuscular model. Prior to the main experiment, the participant performed 15-20 training trials to familiarize the task. After the main experiment, the maximum voluntary contraction (MVC) was recorded in flexor and extensor by asking the participant to maximally contract the respective muscle for 2 s with 5 repetitions. The whole experiment took about 2 hours.

#### Unloading

In the “unloading” session (Fig. 2A), five levels of torque were applied: 0.4, 0.6, 0.8, 1.0 and 1.2 Nm, for flexor (extension torque) and extensor (flexion torque) separately. For each muscle, 10 trials were repeated for each torque level and trials with different torque levels were randomized. Unloading of flexor and extensor was done sequentially. Thus, the unloading session consisted of 100 trials in total. At the beginning of each trial, the participant stabilized the wrist in the position of about 20° flexion against the extensor load or about 20° extension against the flexor load. About 3 s after the initial position was established, the load was abruptly removed (motor turned off). Participants were instructed not to voluntarily correct the hand excursion elicited by unloading and let the hand come naturally to a new position. The unloading task is easy to get familiar with, as when a person holds a heavy book vertically in the palm of one hand and if the book is suddenly lifted off by someone else, the arm moves smoothly to a higher position. The unloading experiment is thus a horizontal version of such a task. 15-20 training trials before the main experiment were sufficient for participants to produce systematic responses to different levels of loads.

**Figure 2.**
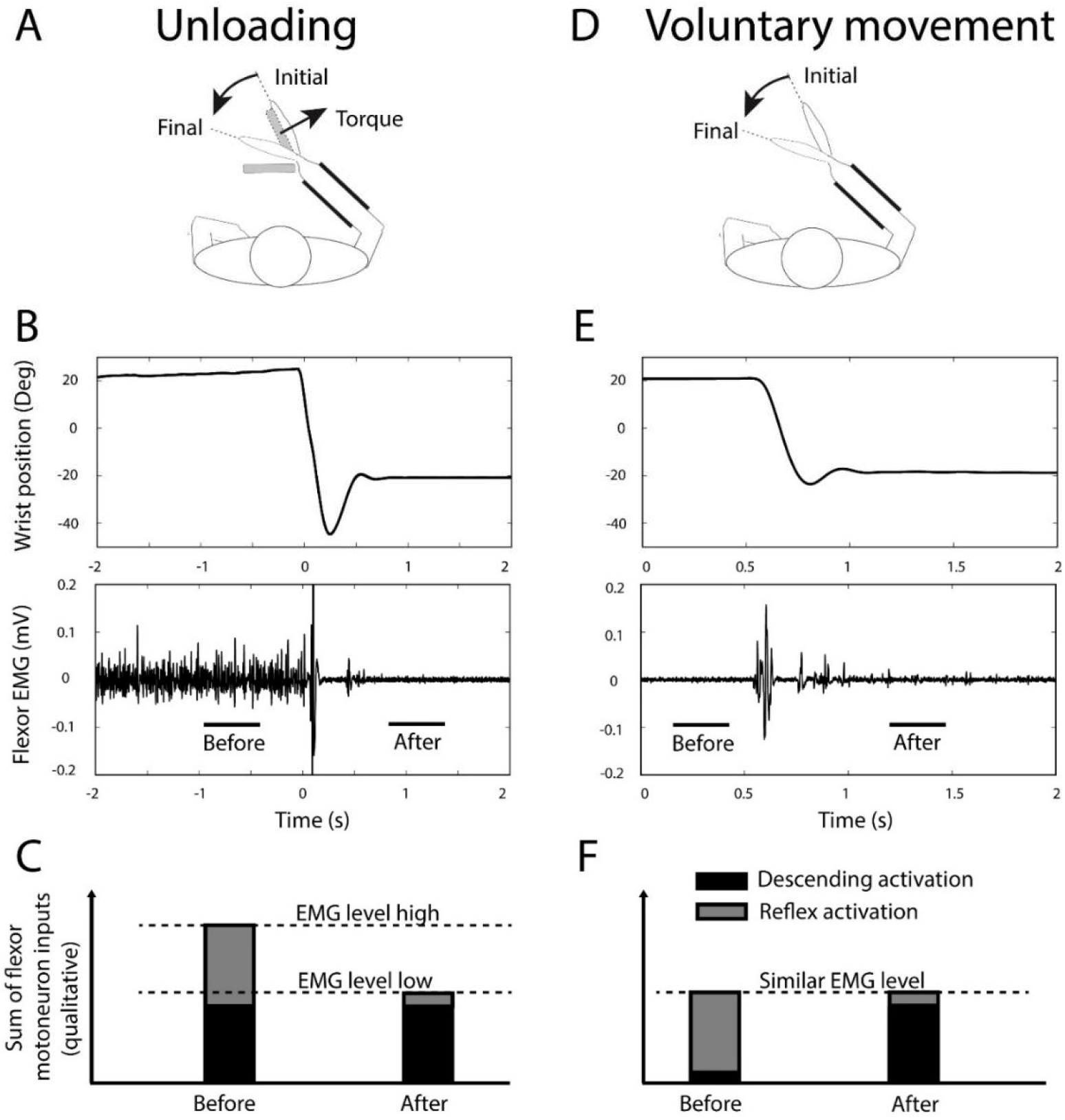
Experimental setup, sample trials of unloading and voluntary movements, and qualitative illustration of descending and reflex activations before and after these movements. (A) Experimental setup of unloading of wrist flexor. In the initial position, the participant was counteracting a constant external torque. The participant was instructed not to voluntarily react to unloading. When the torque load was suddenly removed, the preloaded wrist went to the final position. (B) The time profiles of wrist angle position and raw wrist flexor muscle EMG during unloading. Before unloading, the participant needs to activate flexor muscle to compensate for the load. Time zero represents unloading onset. (C) Qualitative representation to explain the contributions of descending and reflex activations as two major inputs in composing the motoneuron activation before and after unloading, in a wrist flexor muscle. It is assumed that the participant maintained the same descending activations after unloading. After unloading, flexor muscle shortens and the reflex activation decreased, resulting in a lower flexor EMG. (D) Voluntary reaching was flexion from initial to final positions, without any load on wrist. (E) The time profile of wrist angle position and raw wrist flexor muscle EMG during voluntary reaching. (F) Qualitative representation to explain the contributions of descending and reflex activations as two major inputs in composing the flexor motoneuron activation before and after voluntary movement. Similar as the case of unloading, the flexor reflex activation decreased after movement since flexor muscle shortens. After movement, descending activations need to increase to maintain the similar EMG level as before.

#### Voluntary movements

In the session of “voluntary movements” (Fig.2D), wrist reaching movements of both flexion and extension were performed with fast and slow speeds, with 10 trials each. The torque motor was off through this session. These separate sets of movements were performed sequentially for each speed and each muscle, with 40 trials in total. Participants were instructed to always make a single and smooth movement to the target, both for fast (“move as fast as possible”) and slow (“move with a moderate speed”) conditions. In each trial the participant initiated a 40° reaching movement (flexion from 20° to −20, or extension from −20° to 20°) in response to an auditory “go” signal. After each trial, the trajectory of wrist position was displayed on the screen and examined visually by the experimenter. If the movement time fell out of the required range (fast: 0.1-0.3 s, slow: 0.6-0.9 s) for each speed, the participant was informed that the trial was either too slow or too fast, and the trial was repeated.

### Data processing

Wrist angular position data was smoothed by a 40 ms window zero-phase moving-average filter. EMG signals were filtered by a zero-phase 4th order Butterworth band-pass filter (10–500 Hz). These signals were then rectified and low-pass filtered (30 Hz cutoff frequency). Movement onset and offset were defined as the moments when velocity first exceeds or drops below 5% of its peak value, respectively. For group comparison, EMG signals were normalized to percentage of MVC and movement times were normalized to 0-100%.

### Statistics

Values are reported with mean and standard deviation in text and figures. Comparison between conditions in individual participants was done with unpaired t test while group comparison was done with paired t test.

### Reconstruction procedure

The reconstruction procedure was based on a neuromuscular model in which the activation of alpha motoneurons (*A*) depends on descending activation (*u*_*d*_) and the length- and velocity-dependent afferent activations:

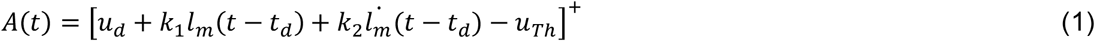

Here, [ ]^+^ is a semi-linear threshold function. The parameters, *k*_1_ and *k*_2_, reflect the sensitivity of muscle spindles to muscle length *l_m_* and its rate of change 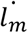. The parameter, *u*_*Th*_, represents the intrinsic threshold of alpha motoneurons. The variable *A* represents the superposition of activation of all alpha motoneurons of a muscle. The variable *u*_*d*_ represents the ensemble of descending activation to these alpha motoneurons. The reflex delay is set as *t*_*d*_ = 20*ms* in wrist muscles.

The recorded muscle EMG is approximated as a linear function of *A*:

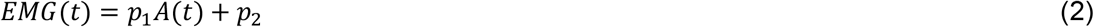

with parameters, *p*_1_,a scaling factor, and *p*_2_,the baseline EMG level when the muscle is fully relaxed.

The combined length of the muscle and tendon, (*l*_*m*_ + *l*_*t*_), is determined by the associated joint angle, *θ*, and approximated as a linear function with a constant moment arm, *c*_1_:

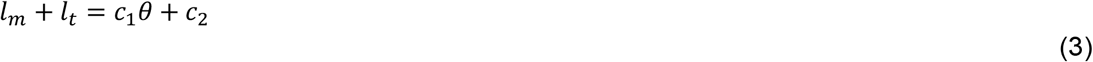

#### Unloading

The analysis of the unloading condition is based on intervals before and after unloading, during which the system is stationary. The reflex delay in Eq. (1) can thus be neglected. Before the onset of unloading, the wrist is approximately stationary 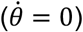 and the muscle is active (see Fig. 2B). The terms 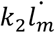 and [ ]^+^ can thus be omitted in Eq. (1). Since participants were instructed not to react to unloading, we assume that descending activation, *u*_*d*_, remains the same *before* and *after* unloading: 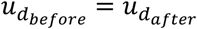 (Asatryan and Feldman 1965, Ilmane et al. 2011).

The tendon force, *F*_*t*_, is approximated as a quadratic function of tendon length, *l*_*t*_ (van Soest and Bobbert 1993):

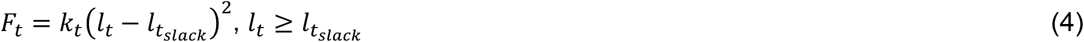

where *k*_*t*_ represents the tendon stiffness and tendon length must be larger than its slack length, *l*_*slack*_. Before the unloading onset, the torque, *T*_*t*_ = *F*_*t*_*c*_1_, produced by the tendon is assumed equal to the force produced by the muscle (equilibrium condition) and counteracts the externally applied torque *T*_*ext*_.

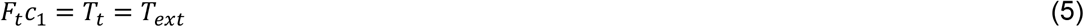

Using equations (1) – (5) both before and after unloading, and introducing parameters *p* = *p*_1_ *k*_1_ *c*_1_ and 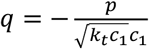 we have (detailed derivation see Appendix - unloading):

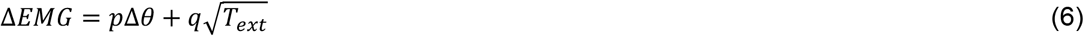

For each participant, a dataset 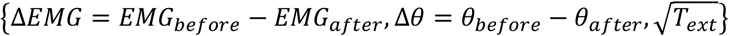 was calculated across load levels *T*_*ext*_, from the mean values across trials of corresponding *EMG* and *θ* data in the interval [−1 −0.5] s *before* unloading onset and [1 1.5] s *after* unloading onset. Postulating that at zero load 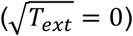, changes in EMG and angles would be zero (Δ*EMG* = 0, Δ*θ* = 0), the 5 load levels, lead to a dataset for regression containing 6 samples. Least-square optimization provided estimates of *p* and *q* with the constraints of *p* ≥ 0 and *q* ≤ 0 in Eq. (6).

The relative contribution of tendon length change to the length change of the muscle-tendon unit, (Δ*l*_*t*)/(Δ*l*_*m* + Δ*l*_*t*) = √(*T*_*ext*/(*k*_*t c*_1))/(*c*_1 Δ*θ*) = (−*q*√(*T*_*ext*))/*p*Δ*θ* was computed from Eqs.3 and 4, and the definitions of *p* and *q* (see also Appendix - unloading). Larger values of this term (as a percentage) indicate that unloading is associated with larger tendon and smaller muscle length change.

This procedure was performed separately for flexor and extensor unloading data. The data of one block (participant No.10, flexor in flexor unloading) out of 20 blocks [2 (conditions of flexor or extensor unloading)*10 (participant number)] was not analyzed because the participant’s EMG across different loads was not separable.

#### Voluntary movements

The relationship (6) between EMG and joint angles can now be used to estimate descending activation from the reflex model (1) during voluntary movements. Rather than directly mapping EMG onto motor neuron activation (equation 2), we use the estimated parameters, *p* and *q*, in Eq. 6 to estimate descending activation in the units of EMG,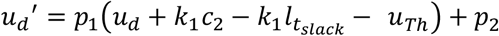. This is possible because during voluntary movement, the muscle EMG values always stayed above threshold, so that the semi-linear threshold function can be resolved. We find (detailed derivation see Appendix – voluntary movements):

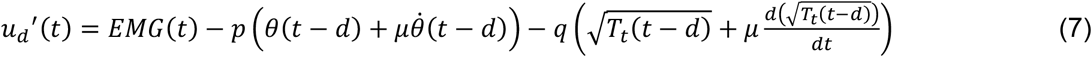

where 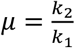. The linear rescaling, *u*_*d*_^′^, of *u*_*d*_ (*t*) preserves its temporal structure, which this study is aimed to uncover. The rescaling also makes descending activation directly comparable with measured EMG (see Results). In fact, from Eq. (7), EMG can be decomposed into a descending, *u*_*d*_^′^(*t*), and a reflex contribution, 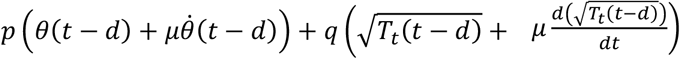.

On the right-hand side of Eq. (7), the values of *p* and *q* have been estimated from the unloading condition, and the variables, *EMG*(*t*), *θ*(*t* − *t*_d_) and *θ*(*t* − *t*_d_) are available as measured data. The parameter *μ* represents the relative weight of length and velocity-dependent components of the afferent signal and is unknown. It In previous studies (Zhang et al. 2016, Hummert et al. 2024) it was set to 0.06s. Different values will be in the present study. The variable, *T*_t_(*t*), represents the torque produced by muscle-tendon forces. Its values cannot be directly computed for each muscle. To provide an estimate we first compute the joint torque *T*(*t*):

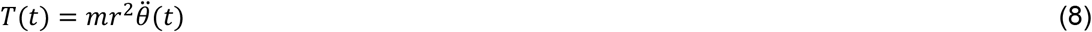

where hand mass, *m*, and radius of wrist flexion/extension, *r*, were taken from standard literature (Winter 2009) as *m* = 0.5 kg and *r* = 0.1 meter. We further assumed that co-contraction was zero in the voluntary movement condition so that only one, either agonist or antagonist, produces torque. For the agonist muscle we have:

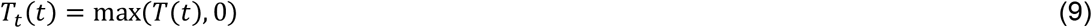

Thus *u*_*d*_*′*(*t*) for the agonist muscle can be obtained in Eq. (7). A similar procedure was performed separately for the antagonist muscle.

For the visual inspection of the time courses of descending, reflex and EMG activations, we focused on the changes these variables experience over the course of the movement. Each function was shifted, therefore, such that its initial value was zero. All three functions are thus plotted relative to this shared baseline. In plots of movement kinematics, the joint angle was also shifted to start at 0.

#### Sensitivity analysis

To establish how sensitive the estimated time courses are to the parameter *μ* we varied its values between 0 and 0.12s.

We similarly performed a sensitivity analysis of the estimated time courses to the level of co-contraction of agonist and antagonist muscles. Co-contraction is defined as a constant contribution to torque by both muscles expressed as a percentage *k* of the peak joint torque observed during voluntary movements. At a given level of co-contraction, *k*, the torque generated by the agonist muscle is:

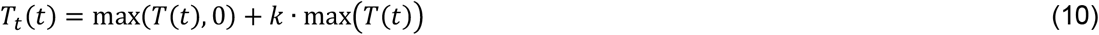

## Results

### Unloading and model calibration

The unloading paradigm is illustrated in Fig. 2B, by one typical trial. At the beginning of the trial, the participant established a wrist extension position in which the flexor muscle was activated to counteract an external load. After the preloaded wrist flexor was abruptly unloaded (at time 0), a new wrist position was established, and the muscle EMG decreased to almost zero (Fig. 2B), significantly smaller than the EMG level before unloading (p<0.01).

Wrist angle positions and EMGs at different load levels are illustrated Fig. 3A for flexor unloading and in Fig. 4A for extensor unloading (data from Participant 4 and 9, respectively). The larger the initial load, the larger the EMG before unloading and the larger the displacement after unloading (blue curves compared to curves in other colors, Figs. 3A and 4A) is the pattern observed across participants.

**Figure 3.**
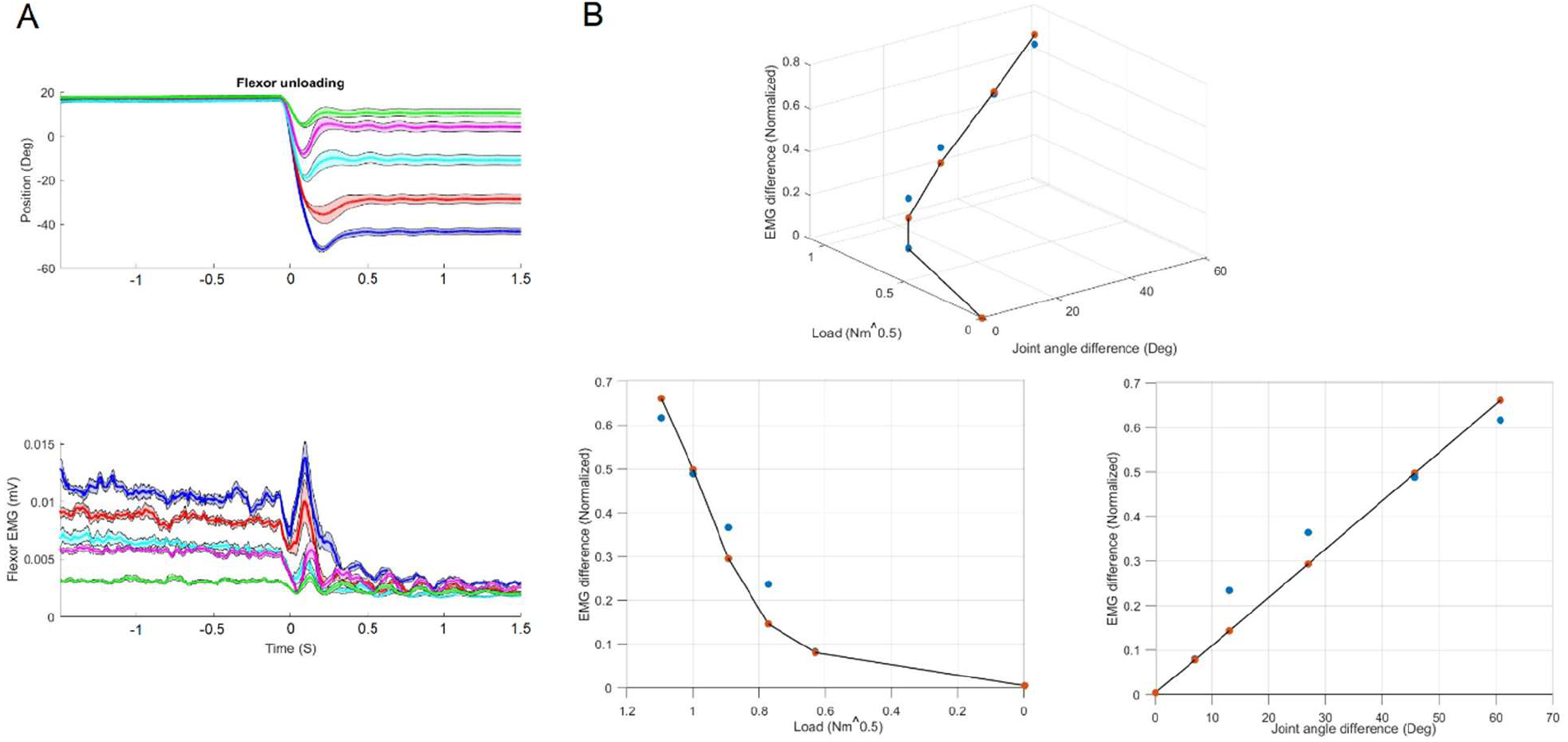
Unloading with different flexor loads in a sample participant (participant No.4). (A) Upper plot: Time courses of wrist joint angle (“position”) at different levels of flexor loads (from 0.4Nm in green to 1.2Nm in blue, mean with standard deviation as band). Lower plot: Time courses of rectified flexor EMG at different flexor loads. Unloading onset is at time 0. (B) Linear regression (black line) of EMG difference (*EMG*_*before*_ − *EMG*_*after*_) as a function of joint angle difference (*θ* _*before*_ − *θ* _*after*_) and the square root of external load 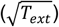 (Eq. 6), plotted together with individual values at each load level (experimental data: blue dots; fitted data: red dots

**Figure 4.**
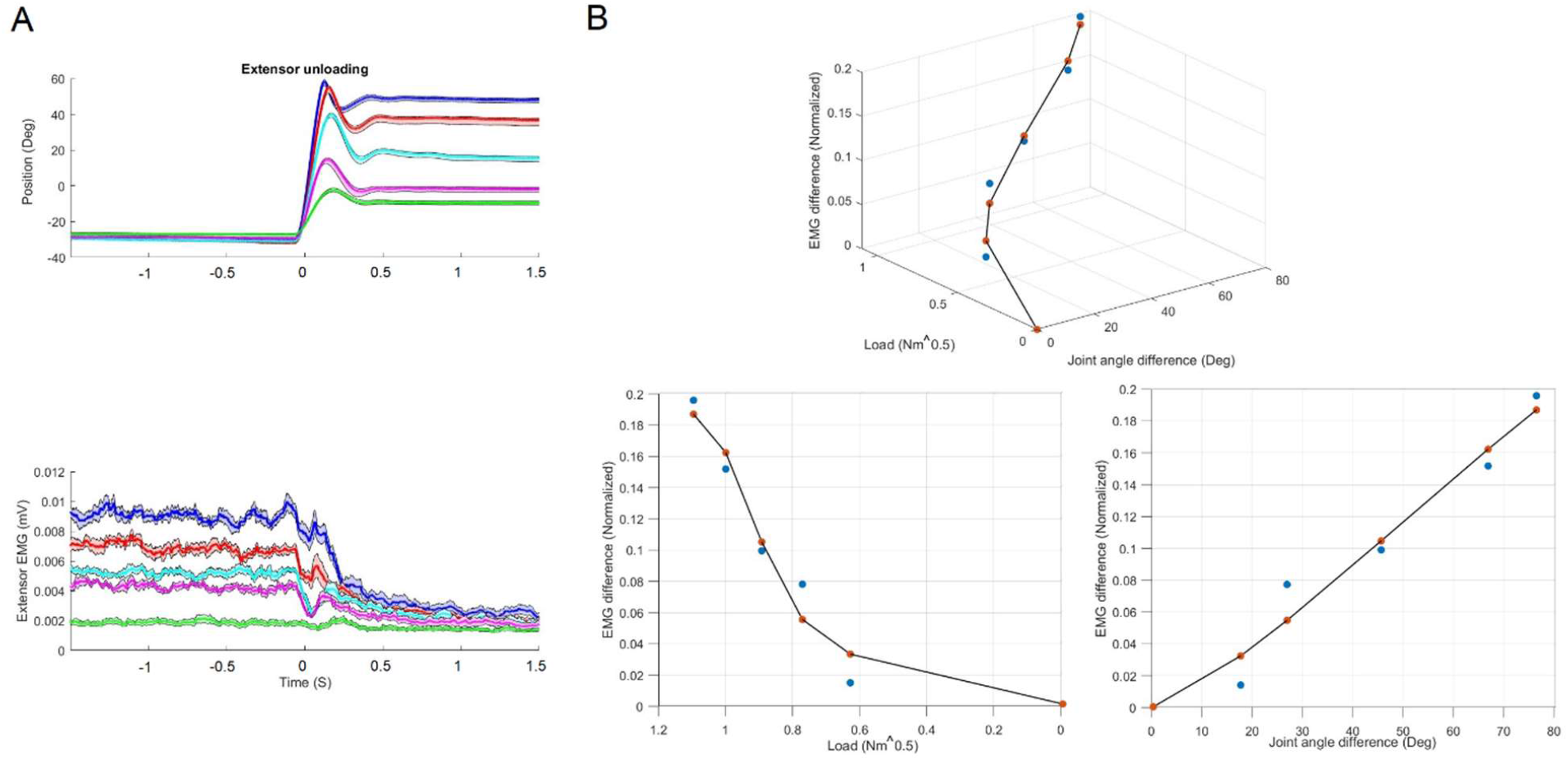
Unloading with different extensor loads in a sample participant (No.9). Same conventions as in Figure 3.

According to Eq. 6, the EMG difference before and after unloading (Δ*EMG* = *EMG*_*before*_ − *EMG*_*after*_) can be modelled as a linear function of joint angle difference (Δ*θ* = *θ*_*before*_ − *θ*_*after*_) and the square root of external load 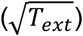. In the first sample participant (Fig.3), the EMG difference could be explained by joint angle difference alone (regression coefficients *p* = 0.011, *q* = 0, *R*^2^ = 0.95, Fig.3B). In the second sample participant (Fig.4), the combination of joint angle difference and the square root of load predicts the EMG difference (regression coefficients *p* = 0.015, *q* = −0.459, *R*^2^ = 0.91, Fig.4B). Note that in this participant, there was no obvious EMG difference at the smallest load (green curves, Fig. 4A), while the joint angle difference was significant.

Table 1 reports the parameter values obtained from the regression of Eq. 6 for all participants. We also report the ratio, Δ*l*_*t*_/(Δ*l*_*m*_ + Δ*l*_*t*_), between tendon length change and length change of the muscle-tendon unit. This ratio predicts the extent to which EMG differences are explained by joint angle differences alone (value near zero) vs. by both joint angle differences and torque (non-zero values). Overall, tendon length varies less in flexor than in extensor unloading (*q* = 0 in 5 out of 9 cases in flexor unloading and in 2 out of 10 cases in extensor unloading).

**Table 1.** Model parameters *p* and *q* obtained from regressing data from the unloading experiment with Eq. 6 for all participants, with goodness of fit reported as *R*^2^. Here, Δ*l*_*t*_ is the tendon length change and Δ*l*_*m*_ is the muscle length change induced by unloading.

| Subj. No. | Flexor |  |  |  | Extensor |  |  |  |
| --- | --- | --- | --- | --- | --- | --- | --- | --- |
| | $p$ | $q$ | $R^2$ | $\frac{\Delta l_t}{\Delta l_m + \Delta l_t}$ | $p$ | $q$ | $R^2$ | $\frac{\Delta l_t}{\Delta l_m + \Delta l_t}$ |
| 1 | 0.0012 | 0 | 0.6111 | 0 | 0.0062 | -0.0137 | 0.9735 | 0.0921 |
| 2 | 0.0084 | -0.2058 | 0.8266 | 0.4889 | 0.0250 | -0.9735 | 0.8838 | 0.8986 |
| 3 | 0.0014 | 0 | 0.7024 | 0 | 0.0197 | -0.3078 | 0.9369 | 0.5984 |
| 4 | 0.0109 | 0 | 0.9549 | 0 | 0.0202 | 0 | 0.9759 | 0 |
| 5 | 0.0014 | 0 | 0.6728 | 0 | 0.0027 | -0.0990 | 0.8852 | 0.7442 |
| 6 | 0.0035 | 0 | 0.7295 | 0 | 0.0078 | -0.3668 | 0.8627 | 0.7754 |
| 7 | 0.0072 | -0.0428 | 0.8805 | 0.1844 | 0.0174 | -0.4967 | 0.7842 | 0.7811 |
| 8 | 0.0033 | -0.1408 | 0.9367 | 0.7225 | 0.0028 | -0.0281 | 0.9695 | 0.2245 |
| 9 | 0.0037 | -0.0331 | 0.8610 | 0.2255 | 0.0150 | -0.4585 | 0.9103 | 0.6547 |
| 10 | - | - | - | - | 0.0027 | 0 | 0.9178 | 0 |

### The time courses of descending activation during voluntary movements

The movement durations are 215.4 ± 16.6 ms, for fast and 779.1 ± 106.7 ms for slow movements. Peak velocities are 363.2±50.9 deg/s for fast and 94.8±18.3 deg/s for slow movements. Movement kinematics and EMG patterns are shown for a sample participant (No.1) in Fig 5 for wrist flexion and in Fig. 6 for wrist extension. The velocity profiles are bell-shaped for both fast and slow movements. For fast, but not for slow movements, the velocity profile often overshoots zero at the end of the movement. In fast movements, muscle EMGs typically begin with an agonist burst followed by an antagonist burst. In slow movements, those bursts are much smaller or even undetectable.

**Figure 5.**
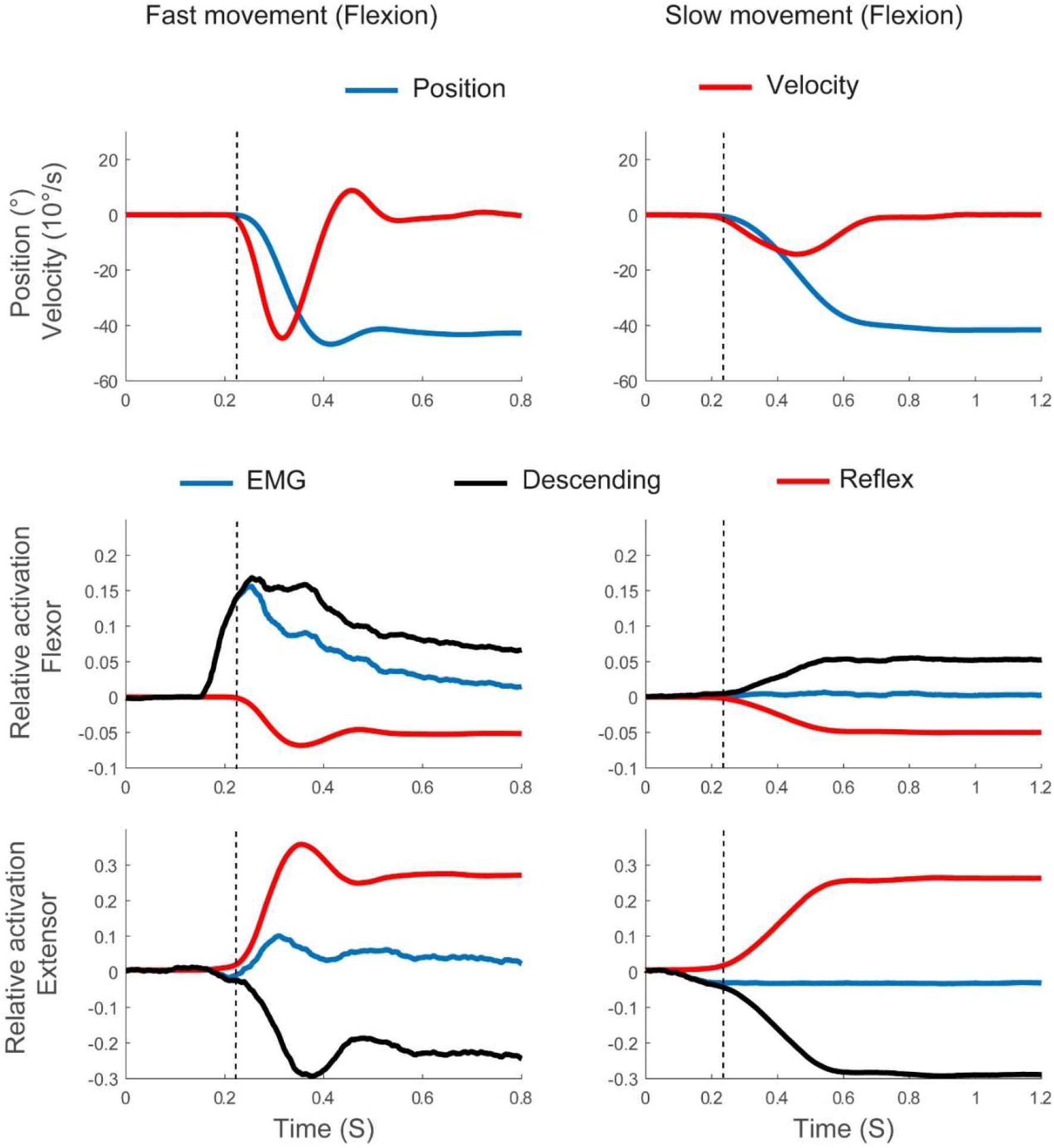
Voluntary wrist flexion movements at two speeds (fast left column; slow right column) for one sample participant (No.1), averaged across trials. Top row plots joint position and velocity. Middle and bottom rows plot reconstructed descending activation together with EMG and reflex activation, in agonist and antagonist muscles respectively. Vertical dashed line indicates movement onset. To enable visual comparison of the time courses, EMG, descending and reflex activations are shifted to all start at 0.

**Figure 6.**
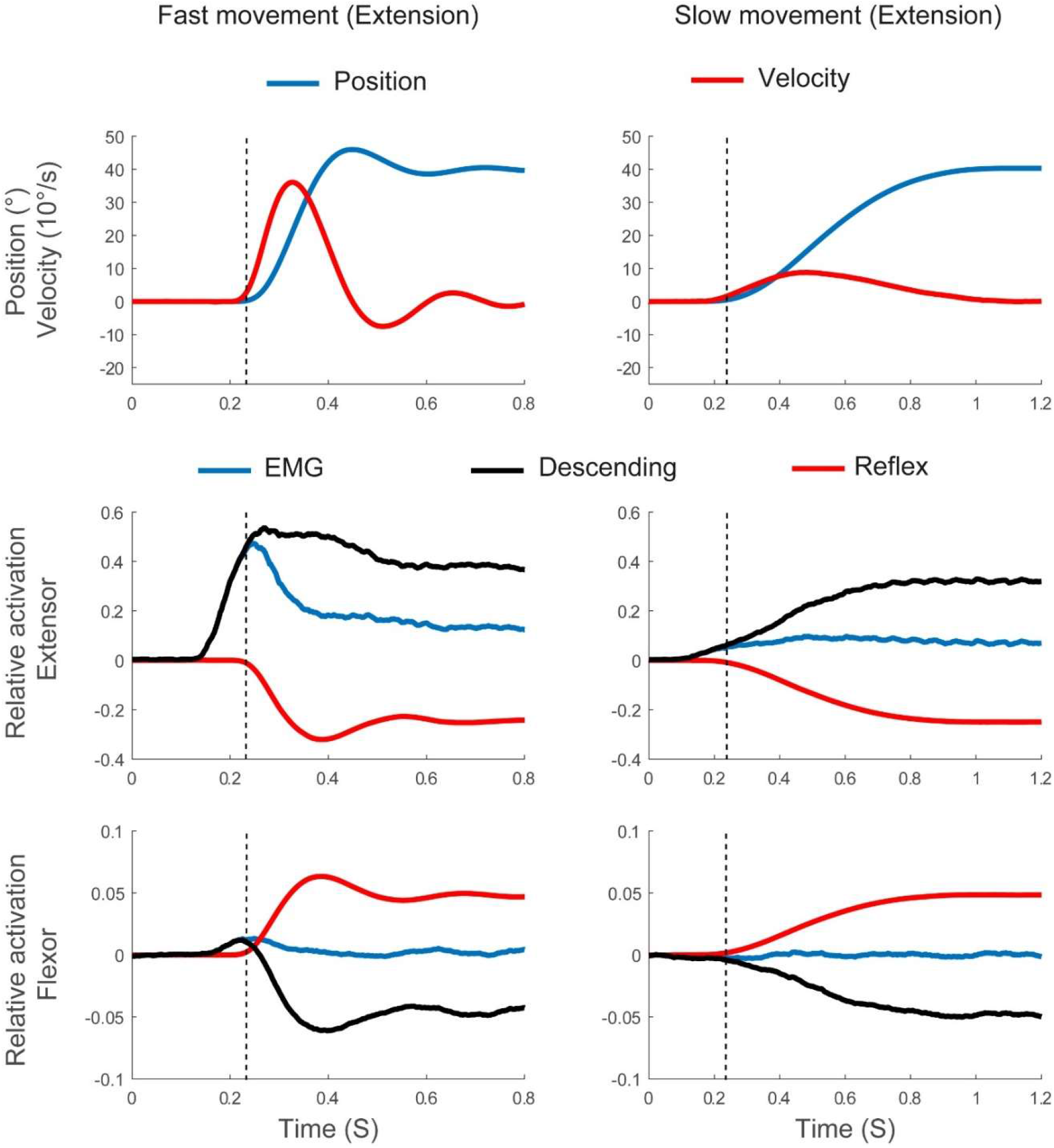
Voluntary wrist extension movements at two speeds (fast: left column; slow: right column) for the same participant as in Fig.5, averaged across trials. Same plotting conventions as in Fig. 5

The time course of descending activation (black curves) has a strong tonic component for both agonist and antagonist muscles. Thus, the level of descending activation at the end of a movement clearly differs from the initial level, while EMG tends to return toward its initial level after a phasic activation burst. Descending activation arises about 100ms before the movement onset. In this first phase, EMG is aligned with descending activation. EMG begins to deviate from descending activation after movement onset, when the reflex begins to modulate EMG. For agonists, the reflex contribution lowers EMG, and, for antagonists, the reflex contribution boosts EMG. The time course of descending activation is ramp-like for slow movements, while it contains a non-monotonic component early in the movement for fast movements. Group averaged patterns of descending activation shown in Figs. 7 and 8 confirm these qualitative features.

**Figure 7.**
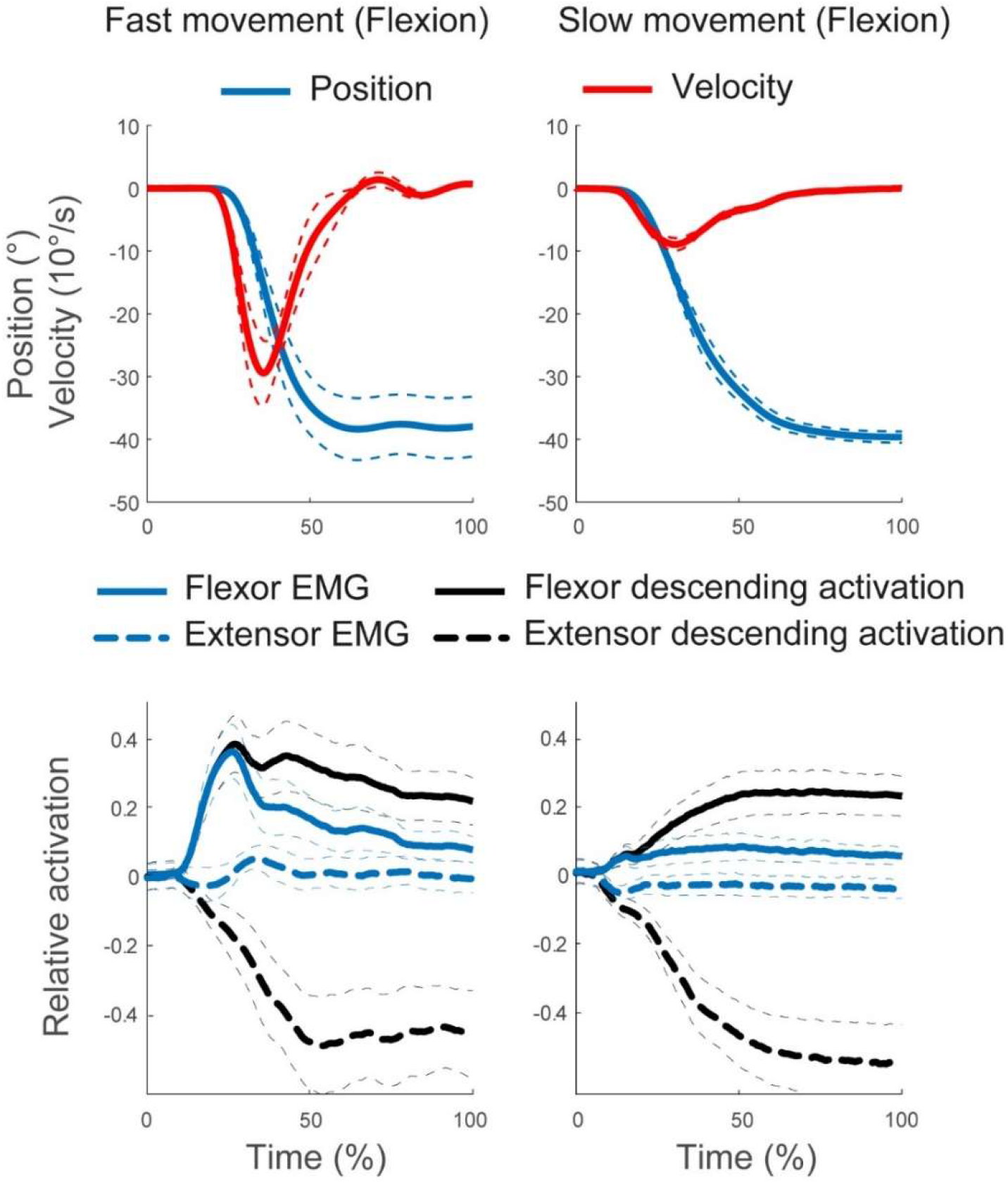
Group average of movement kinematics, EMG, and descending activation patterns during voluntary wrist flexion movements at two speeds (fast: left; slow: right). Movement times are normalized. To enable visual comparison of the time course, EMG and descending activation patterns are shifted to start at 0.

**Figure 8.**
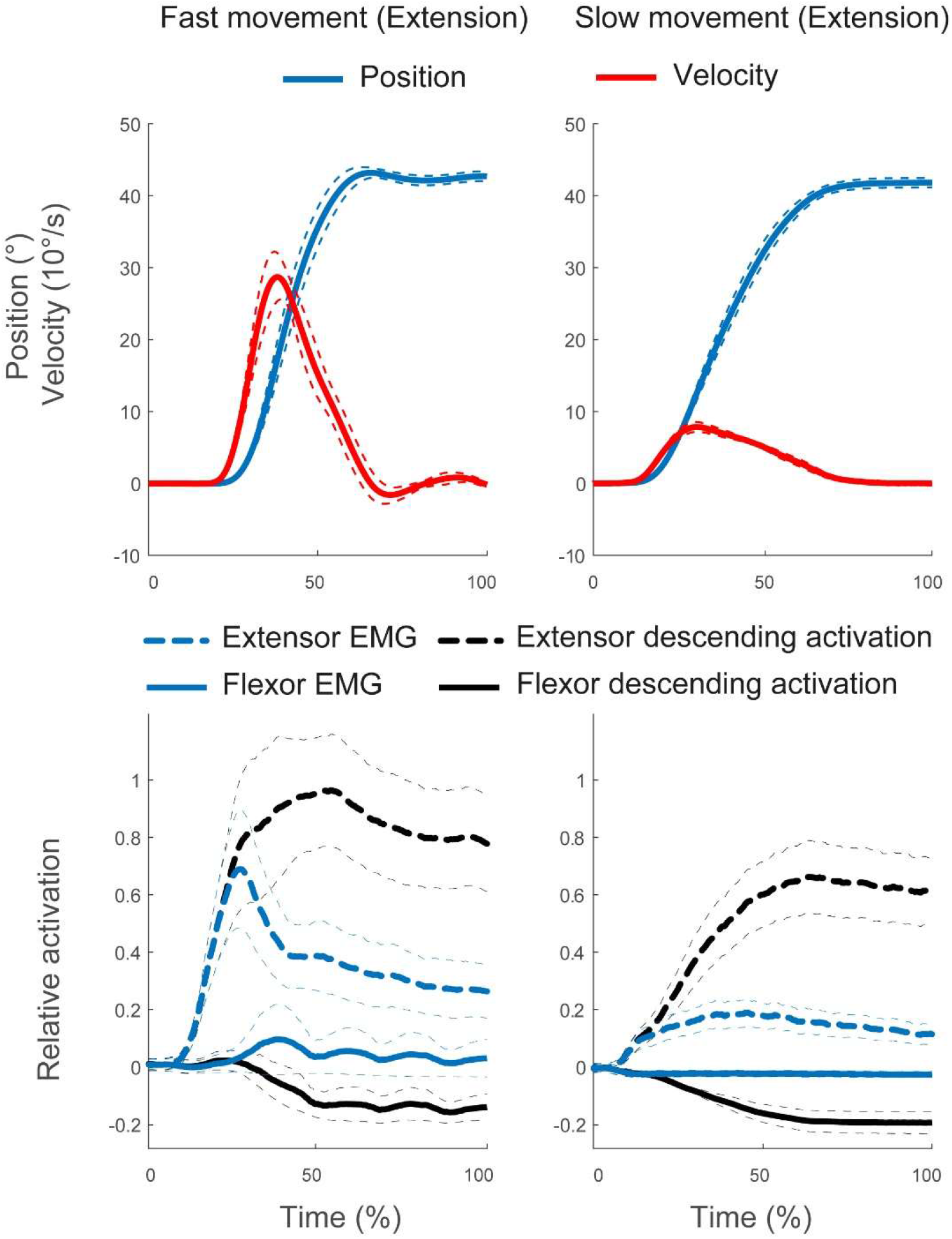
Group average of movement kinematics, EMG, and descending activation patterns during voluntary wrist extension movements at two speeds (fast: left; slow: right). Same plotting conventions as in Fig. 7.

### Sensitivity analysis of descending activation to reflex velocity gain and level of co-contraction

Figure 9 illustrates how strongly the estimated time course of descending activation depends on the gain, 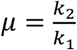, of the velocity contribution to the stretch reflex relative to the position contribution (Eq. 1). This gain cannot be estimated from the unloading data. We explored the range of values from 0 to 0.12s. The velocity component of the stretch reflex only plays a role after movement onset once considerable muscle length changes set in. As a result, the early portion of the time course of descending activation is unaffected by this parameter. During the movement, the modulation of descending activation is moderate for fast and negligible for slow movements.

**Figure 9.**
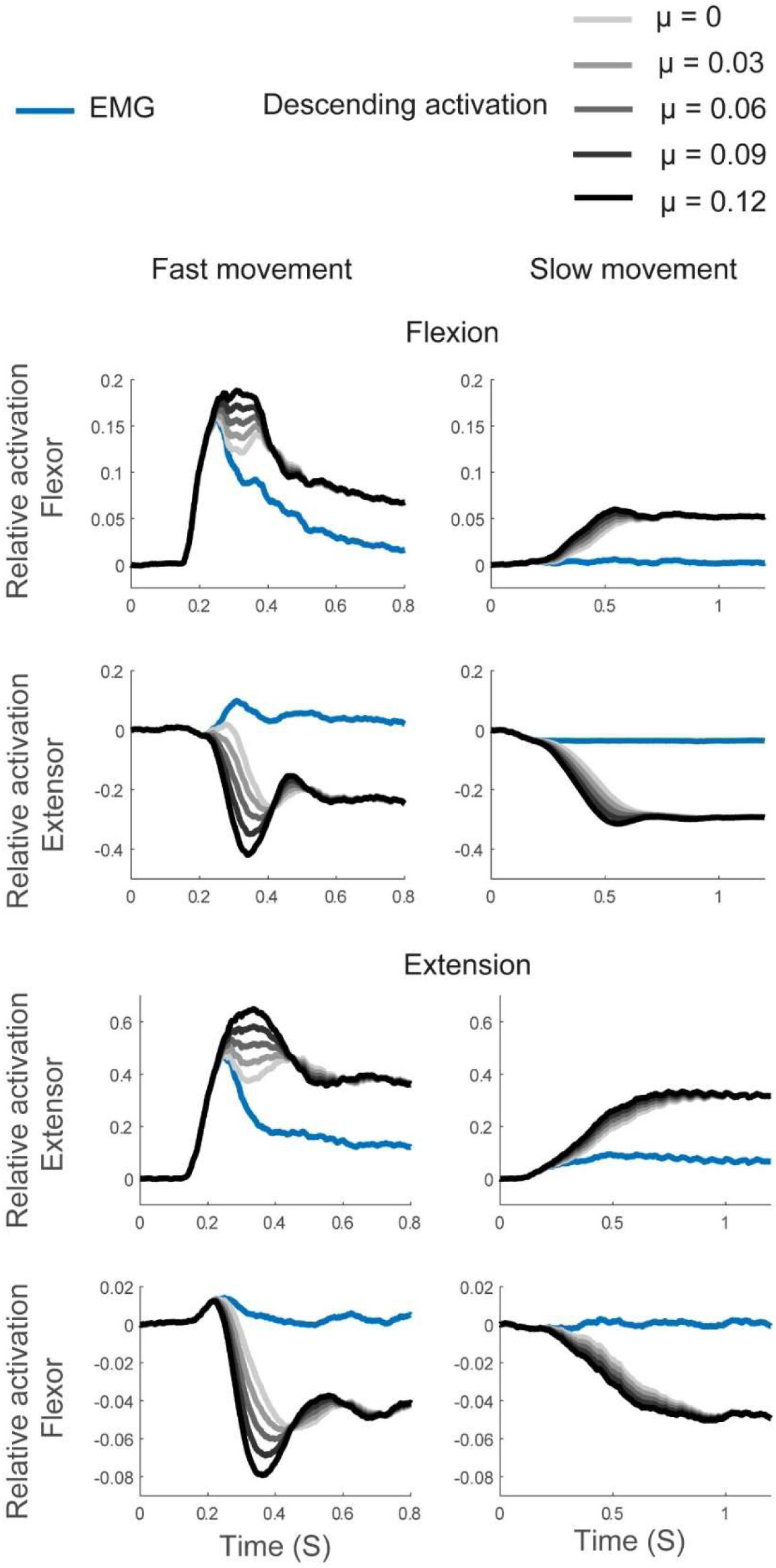
Descending activation estimated at varied levels of *μ* ∈ [0, 0.12]*s* for fast (left column) and slow movements (right column), in a sample participant (No.9). To enable visual comparison of their time courses, EMG and descending activation at *μ* = 0.06 were shifted to start at 0. Descending activation at other *μ* values were shifted together with *μ* = 0.06, maintaining their relative vertical positions.

Figure 10 illustrates how strongly the estimated time course of descending activation depends on co-contraction. Co-contraction cannot be directly estimated from the observed EMG, as that would require calibrating EMG to actual levels of force. We instead varied assumed levels of co-contraction, expressed as a percentage of maximum joint torque, ranging from 0 to 100%. Clearly, even this broad range of co-contraction levels does not substantially change the time structure of descending activation.

**Figure 10.**
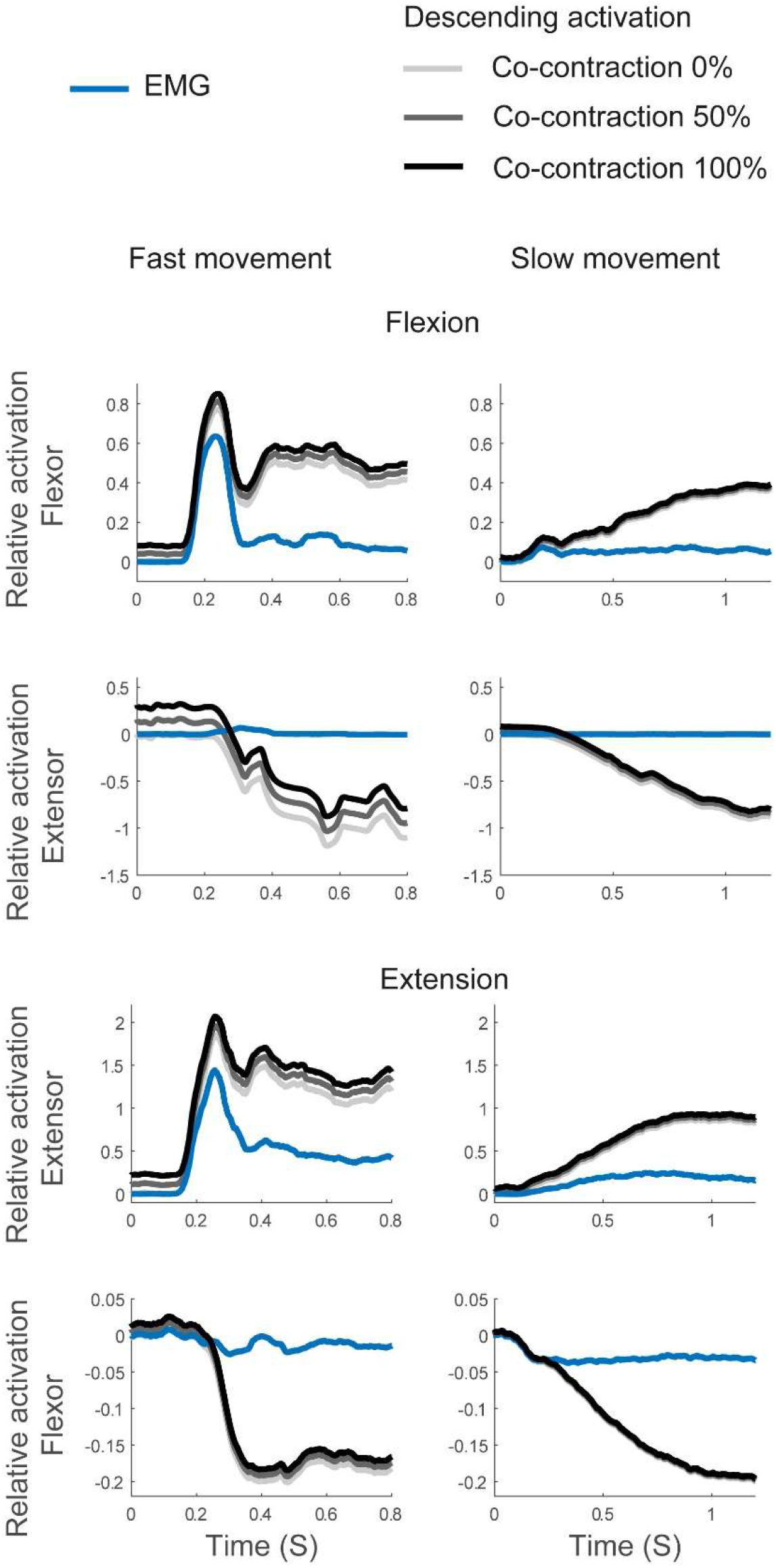
Descending activation estimated at varied levels of co-contraction (0-100% of peak joint torque), for fast (left column) and slow movements (right column), in a sample participant (No.2). To enable visual comparison of their time courses, EMG and descending activation at 0% co-contraction were both shifted to start at 0 while descending activation at 50% and 100% were shifted together with the estimate at 0% co-contraction, maintaining their relative vertical positions.

## Discussion

This study used a model of the spinal stretch reflex to estimate patterns of descending activation from measured EMG and kinematic time courses of wrist flexion and extension. This enabled us to observe three key features of the descending activation. (1) Muscle activation is aligned with descending activation initially, but after movement onset the time courses of descending and muscle activation differ qualitatively, reflecting reflex contributions to muscle activation. (2) The time course of descending activation is monotonic for slow movements, leading from an initial level to a final level that is shifted during the movement. For fast movements, descending activation reaches the same final level, but evolves non-monotonically during the movement. (3) Different levels of the gain of the velocity-dependent component of spinal feedback and of co-contraction do not lead to qualitatively different time courses of estimated descending activation.

### How descending and reflex activation jointly shapes muscle activation

Our model-based estimates demonstrate that the spinal reflex requires that descending and muscle activations have qualitatively different time courses. The two are naturally aligned before movement onset, that is, before sensory feedback signals have begun to change (Figs. 5 and 6). As movement starts, agonist muscles shorten and antagonist muscles lengthen so that the spinal reflex begins to contribute to muscle activation. The separation of descending and muscle activations continues throughout the movement. At the end of the movement muscle activation returns to its initial low level (except for changes in co-contraction), while agonist descending activation settles at a higher than its initial level. This compensates for the reduced reflex signal from the shorter agonist muscle. Antagonist descending activation settles at a lower than its initial level. This compensates for the enhanced reflex signal from the now longer antagonist muscle and prevents the stretch reflex from activating the antagonist muscle (Figs.5-8 and Fig.2F). As a result, descending activation has strong tonic component that reflects the need to adjust the effective reflex threshold to the shift of posture that results from a voluntary movement (Feldman and Levin 1995, Albert et al. 2020).

### Descending activation patterns are reshaped to move at different speeds

Is shifting the threshold of the stretch reflex alone sufficient to achieve voluntary movement at a given speed? We saw that movement is initiated by directly driving muscle activation by descending activation. This could be viewed as the initial phase of a shift in reflex threshold that continues throughout the movement (Feldman 1986). In this perspective, different movement speed results from different rates of this shift, with potential differences in the timing of these shifts for agonist vs antagonist muscles that modulate co-contraction (Feldman and Latash 2005). That is one interpretation of earlier controversial discussions on the complexity of the time courses of descending activation (Gomi and Kawato 1996, Gribble et al. 1998, Kistemaker et al. 2006). We show that estimated patterns of descending activation at high speeds are not mere rescalings of the patterns estimated at low speeds. Instead, the time course of descending activation at high speeds is more complex than the time course at low speeds, with a small non-monotonic component, “phasic-tonic” pattern. Increasing co-contraction up to 100% of peak joint torque did not change this “phasic-tonic” pattern. This difference in the time structure of descending activation at higher speeds is consistent with similar results obtained from two different methods of estimating descending activation (Ramadan et al. 2022; Hummert et al. 2024). Both studies found that varying co-contraction did not substantially affect this outcome.

These results do not exclude a general role for the modulation of reflex gain (Azim and Seki 2019) or of co-contraction. Note, however, that the estimates are conservative relative to such modulation: If the reflex was suppressed during movement, it would need to be re-instated at the end of the movement, leading to an even more strongly temporally structured pattern of descending activation. Expanding the approach to include more complete spinal circuits (Windhorst 2007) would strengthen the methodology but requires non-trivial methodological innovation.

### Model-based approaches for estimating descending activations

Many contemporary computational models of voluntary movement generation do not explicitly address the contribution of spinal reflex circuits (Todorov 2000, Todorov and Jordan 2002, Scott 2012, Churchland and Shenoy 2024, see also Weiler et al. 2019&2021). There is, however, a considerable theoretical literature around the role of the spinal reflex and of more complex spinal reflex circuitry (Gribble e al. 1998, Raphael et al. 2010, Zhang et al. 2016, Buhrmann and Paolo 2014, Tsianos et al. 2014, Blum et al. 2020, Niyo et al. 2024). This literature establishes the contributions of reflex circuits to movement generation by inserting different kinds of pre-defined temporal profiles of descending activation and analyzing the resultant movement kinematics or force patterns. In some cases, such pre-defined temporal patterns of descending activation were parametrically varied to find the profile best matching kinematic data (Frenkel-Toledo et al. 2019, Zhang et al. 2022).

The present approach builds on two previous extensions of these paradigms which estimated the time course of descending activation without pre-defined templates. Hummert et al. (2024) employed a model consisting of the standard biomechanics of a planar, two-joint arm, of a relatively simple muscle model (Gribble et al., 1998), and the same reflex model used here. With the help of additional simplifications, this model could be inverted analytically to estimate descending activation patterns for four muscles directly from the kinematics of human movement data obtained at two speeds. The results are qualitatively consistent with the results reported here. Ramadan et al. (2022) employed the same model with six muscles (thus including muscle redundancy). They found time courses of descending activation that change minimally from initial to the final posture, while generating the movement within a given movement time, using numerical methods from optimal control. The resultant patterns are constrained only by movement start and endpoint and movement time, not containing any fit parameters. This study too found the same qualitative time structure of descending activation as in the present study.

In the present approach, descending activation can be estimated directly from measured EMG patterns, without the need for a muscle model. This eliminates any dependency of the estimated descending activation on parameters of the muscle model. A key step of this approach is to use unloading experiments to estimate the critical model parameter (*p* in Eq. 6) that characterizes the relationship between joint angle and EMG change. The contribution of the tendon to this relationship that varies across participants (Table 1) is taken into account. The linear regression model (Eq. 6) captured the observed the relationship between changes in joint angle and EMG (*R*^2^ >0.61 across all conditions). Further model parameters that merely linearly rescale the estimates of descending activation could be eliminated, simplifying the estimation method.

An earlier attempt to estimate descending activation directly from EMG patterns was made in Latash and Goodman 1994. This approached used pre-defined temporal profiles of descending activation together with a similar reflex model to estimate parameters that defined the templates. The templates found to account for observed EMG patterns were phasic-tonic in the same sense as described here.

### Neurophysiological interpretation of the estimated descending activation patterns

By introducing *u*_*d*_′ as a linear function of descending activation *u*_*d*_ (Eq. 7), descending activation was estimated up to unknown scaling parameters, such as the motoneuron threshold. It is therefore the time structure, but not the absolute level of descending activation that can be compared to neural activation patterns from the neurophysiological literature.

When monkeys make load-free reaching movements with an arm, phasic-tonic activation is commonly observed in neurons in primary motor cortex, M1 (e.g., Fig. 12 Kalaska et al. 1989). This pattern is also seen when gravity compensation plays a role (e.g. Fig. 4A, Takei et al. 2018). Recording motor cortical and somatosensory afferent neurons during a reach-to-grasp task, Umeda et al. (2022) observed phasic-tonic patterns in cortical neurons, while afferent activation began immediately after movement onset. They concluded that muscle activation before movement onset was primarily caused by cortical descending input, while activation after movement onset was driven by both descending and afferent inputs. Overall, these neural observations are consistent with the estimates reported here.

Studies observing EMG responses to non-invasive brain stimulation in humans suggest that descending activation to agonists is larger after movement onset than before, while the opposite is true for antagonists (Raptis et al. 2010; Zhang et al. 2017, 2018, 2020). This observation aligns with the theoretical prediction illustrated in Fig.2F. It is also consistent with the pattern observed for the tonic components of the patterns of descending activation estimated before and after movement initiation here (Figs. 5-8).

## Conclusion

Using a simple spinal reflex model to estimate the relative contributions of descending and reflex activations to muscle activation directly from EMG data we obtained insights into the temporal structure of descending activation patterns. Convergent with previous results that were based on different estimation methods, the time structure of descending activation was found to be qualitatively different from muscle activation, containing both tonic and phasic components that vary with the speed of voluntary movement beyond a mere temporal rescaling. These patterns are broadly consistent with neurophysiological findings.

## Acknowledgements

We thank Anatol Feldman for allowing us to use his experimental setup and for general support. We thank John Kalaska for very helpful discussions of the neurophysiological literature. This project has received funding from the European Union’s Horizon 2020 research and innovation programme under the Marie Sklodowska-Curie grant agreement No. 956003.

## Appendix derivation of equations in reconstructing the descending activation patterns

### Unloading

For both before and after unloading, taken together equations (1–3):

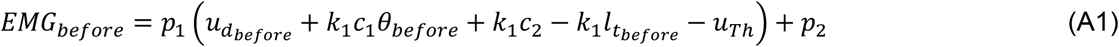

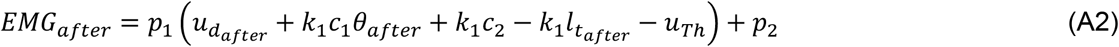

Then subtracting the two equations (A1) and (A2), and defining *p* = *p*_1_*k*_1_*c*_1_:

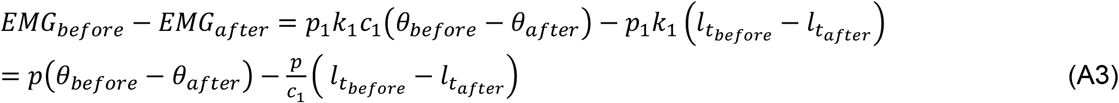

Taking (4) and (5) we can calculate the tendon length before unloading onset:

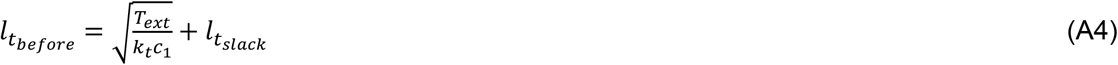

After unloading we have:

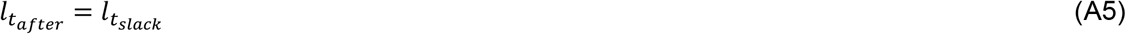

Taking (A3), (A4) and (A5) and defining 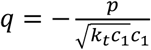:

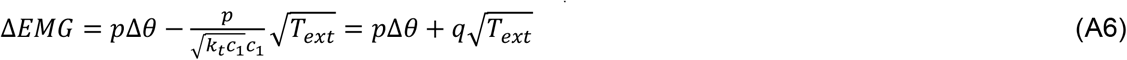

#### Voluntary movements

Taking equations (1)-(3) and considering the reflex delay *d* we have:

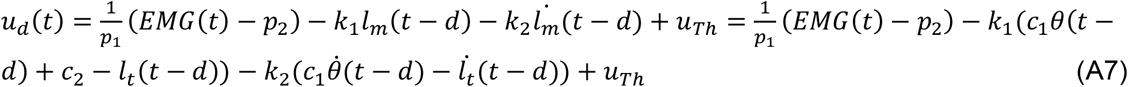

From equations (4) and (5) we have:

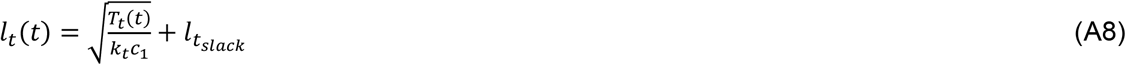

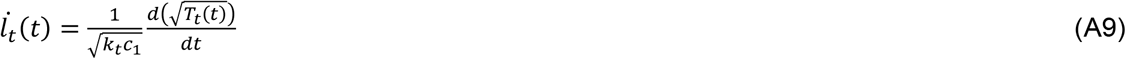

Taken equations (A7) - (A9), and let 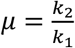:

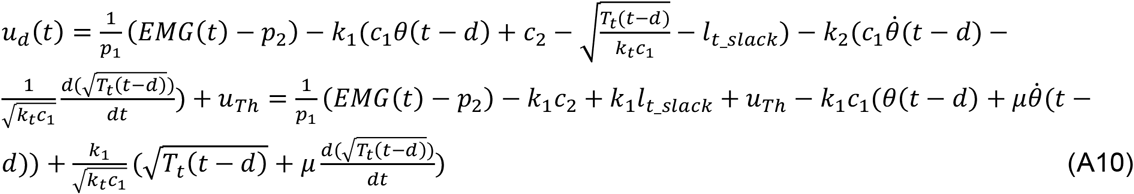

Reform equation (A10):

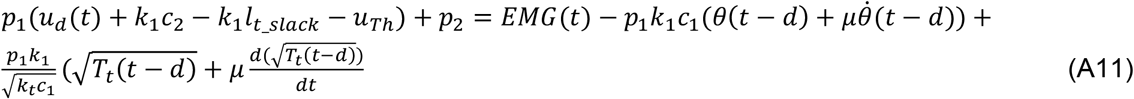

Note that 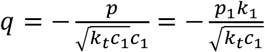 and let *u*_*d*_*′* = *p*_1_(*u*_d_ + *k*_1_ *c*_2_ − *k*_1_ *l*_*d_slack*_− *u*_*Th*_)+ *p*_2_ we have:

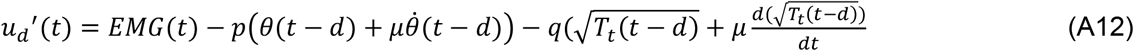

## Notes

### Competing Interest Statement

The authors have declared no competing interest.

